# Assessing Technical College Students’ Biology Proficiency as Related to Gender and Performance in Biology in Nigeria

**DOI:** 10.1101/2024.10.28.620585

**Authors:** Kehinde Sowunmi, Ajoa Iyanu Enoch, Ogbonnaya Uchenna

## Abstract

This study investigated biology proficiency in relation to gender and performance among 350 Nigerian technical college students from 5 elitist technical colleges in Lagos State, employing a quantitative research method within a descriptive survey design. Data collected were analysed using descriptive statistics of frequency, percentage, mean, and standard deviation, alongside inferential statistics including independent samples t-test and multiple regression analysis. Findings revealed that technical college students from elitist schools displayed a high level of biology proficiency. Significant correlations were found among students conceptual understanding, procedural knowledge, strategic competence, adaptive reasoning, productive disposition, and overall performance in biology. While gender differences in biology proficiency appear to be diminishing across most subscales, subtle gender differences were observed in performance in biology in this study. Gender, conceptual understanding, productive disposition, adaptive reasoning, strategic competence, and procedural knowledge made statistically significant contributions to the variance in technical college students performance in biology. Based on this baseline study, it is recommended that future research in Nigeria should explore the biology proficiency of non-elitist schools, which constitute the majority of educational institutions in the country, to enable broader generalization of these findings.

## INTRODUCTION

Biology, as a foundational science subject, is pivotal in shaping students’ understanding of life processes and natural systems. Its mastery has implications for scientific literacy and opens pathways to careers in health sciences, environmental studies, and biotechnology (Olatoye & Ogunkola, 2022). However, achieving proficiency in biology presents unique challenges, especially in Nigeria, where disparities in educational resources and socio-economic factors affect student outcomes (Adekunle & Salami, 2023). Moreover, gender disparities in science education have long been a subject of global discourse, with research suggesting that social and cultural factors contribute to variations in students’ engagement and performance in science subjects, including biology (Chukwu et al., 2021).

Gender-related variations in science proficiency remain a key area of investigation, particularly in developing countries where educational resources and instructional practices often differ markedly by school type (Akinola et al., 2021). Studies have shown mixed results regarding the influence of gender on biology proficiency. While some studies report negligible gender differences in biological sciences (Ibrahim & Ayo, 2021), others indicate that male and female students may exhibit distinct learning patterns and motivational factors, potentially impacting their academic performance (Usman & Olumide, 2022). Addressing these gender dynamics in biology education is essential, as it can inform policies and practices that foster inclusive and equitable learning environments.

The Nigerian educational landscape is further complicated by the division between elitist and non-elitist schools, with elitist schools often benefiting from better infrastructure, instructional resources, and qualified teachers (Olowo et al., 2023). This division can lead to disparities in academic achievement, as students in elitist schools generally perform better in science subjects, including biology (Eze & Ifeoma, 2022). Yet, studies focusing specifically on biology proficiency within technical colleges are sparse, despite the fact that these institutions serve a significant segment of Nigeria’s youth, preparing them for technical and vocational careers where a strong foundation in biology is crucial (Adeyemi & Bello, 2023).

This study, therefore, seeks to fill this gap by examining biology proficiency among technical college students in Nigeria, with a particular focus on gender and its relation to performance. By employing a quantitative approach within a descriptive survey framework, this study aims to understand not only the level of biology proficiency in elitist technical colleges but also the extent to which gender influences performance. This research is guided by the following objectives: (1) to assess the biology proficiency of technical college students, (2) to examine gender differences in biology performance, and (3) to identify specific factors, such as conceptual understanding and procedural knowledge, that contribute to performance variance among students. The insights gained will contribute to the growing body of knowledge on science education in developing countries and inform strategies for enhancing biology education within Nigerian technical colleges.

In line with previous research on science education, this study hypothesizes that gender may exhibit subtle yet statistically significant effects on biology performance, influenced by underlying competencies such as conceptual understanding and strategic competence (Chukwu & Oluwole, 2022). By focusing on these competencies and examining them through the lens of gender, this study will provide valuable insights into the mechanisms shaping biology achievement in technical education settings and lay the groundwork for future research on non-elitist institutions, thus broadening the generalizability of the findings.

### Purpose of the Study

The present study investigated biology proficiency among Nigerian technical college students, specifically examining differences in proficiency between male and female students and exploring the relationship between biology proficiency and overall performance in biology. The study aims to provide insights into the factors that influence proficiency in biology within technical education settings, contributing to a broader understanding of gender dynamics and performance outcomes in Nigerian science education.

### Research Questions

- Is there a significant difference in biology proficiency between male and female students in technical colleges?
- What factors contribute to higher or lower performance in biology among these students?
- How do technical college students’ biology proficiency levels compare across regions or schools in Nigeria?

## METHOD

This study utilized a quantitative research approach within a descriptive survey design to assess biology proficiency among Nigerian technical college students, with a focus on gender differences and the relationship between proficiency and performance. The target population included senior technical college students from elitist technical colleges in Lagos State, and a multistage sampling method selected 350 students from 5 institutions, ensuring gender balance through stratified random sampling. Data were gathered using two instruments: a Biology Proficiency Test (BPT) and a Student Demographic and Performance Questionnaire (SDPQ). The BPT, aligned with core curriculum standards, evaluated proficiency across conceptual understanding, procedural knowledge, strategic competence, adaptive reasoning, and productive disposition. Expert-reviewed for content validity, the BPT was pilot-tested, achieving a Cronbach’s alpha of 0.82, demonstrating strong reliability. The SDPQ collected demographic information, performance data, and factors such as study habits and access to resources. Data collection, conducted over two weeks during supervised sessions, involved participant consent and confidentiality to ensure ethical compliance. Data analysis included descriptive statistics (frequency, mean, standard deviation) to summarize demographic and performance data, and inferential statistics via independent samples t-test and multiple regression. The t-test assessed gender differences in biology proficiency, while multiple regression evaluated the contribution of each proficiency domain (conceptual understanding, procedural knowledge, etc.) to overall performance in biology. Reliability of the instruments was confirmed, with both tools achieving Cronbach’s alpha coefficients above 0.8, and validity was strengthened through expert reviews and factor analysis. Ethical considerations included anonymity, informed consent, and secure data storage. This rigorous methodology provides a robust basis for examining proficiency and performance in biology, specifically exploring gender effects and proficiency components in Nigerian technical education.

## RESULTS AND DISCUSSION

### Research Question One

Is there a significant difference in biology proficiency between male and female students in technical colleges?

A total biology proficiency score was computed from five dimensions: conceptual understanding, procedural knowledge, strategic competence, adaptive reasoning, and productive disposition. For conceptual understanding, scores ranged from zero to 30, for procedural knowledge, scores ranged from zero to 30, for strategic competence, scores ranged from zero to 20, for adaptive reasoning, scores ranged from zero to 15, and for productive disposition, scores ranged from 15 to 90. Together, the biology proficiency score ranged from 15 to 185, with a midpoint score of approximately 100, indicating that scores above this point reflect high biology proficiency.

Of the 350 technical college students sampled, 345 (98.57%) scored above 100 (M = 128.45, SD = 11.05, score range: 80-160, 95% CI = 127.23–129.67), while 5 students (1.43%) scored exactly 100 (M = 100, SD = 0, score range: 100, 95% CI = 100). These results indicate that a large proportion of the students demonstrated high biology proficiency, with an overall mean score for the entire sample of M = 128.35, SD = 11.10, score range: 80-160, and 95% CI = 127.13– 129.57. This distribution suggests that technical college students in this sample generally exhibit strong biology proficiency.

In Table 1 below, descriptive statistics, including mean and standard deviation values, and t-test values are presented for biology proficiency scores and overall biology performance scores among male and female technical college students. Regarding the biology proficiency score, female students achieved a slightly higher mean score (M = 78.56, SD = 10.12) compared to their male counterparts (M = 77.92, SD = 10.24), although this difference was statistically not significant (t348 = -0.52, p = 0.60). In conceptual understanding, female students scored marginally higher (M = 15.12, SD = 3.04) than male students (M = 14.98, SD = 3.11), with this difference also proving statistically insignificant (t348 = -0.35, p = 0.73). Similarly, in procedural knowledge, the female students’ mean score (M = 16.02, SD = 2.85) was slightly higher than that of the male students (M = 15.89, SD = 2.88), yet again with no statistically significant difference (t348 = -0.32, p = 0.75). However, in strategic competence, male students scored marginally higher (M = 13.40, SD = 3.12) than their female counterparts (M = 13.15, SD = 3.05), but this difference was also non-significant (t348 = 0.81, p = 0.42). For adaptive reasoning and productive disposition, no significant differences were observed across genders, indicating similar proficiency levels in these subscales. Notably, male students exhibited a higher mean score in overall biology performance (M = 68.41, SD = 15.32) than female students (M = 64.29, SD = 14.97), a difference that was statistically significant (t348 = 2.15, p = 0.03), suggesting potential gender-based variations in biology performance.

**Table 1:** Independent Samples T-Test Analysis of Technical College Students’ Performance in Biology and Biological Proficiency According to Gender.

| Gender | N | Mean | SD | df | t | p |
| --- | --- | --- | --- | --- | --- | --- |
| Conceptual Understanding | Male | 175 | 14.98 | 3.11 | 348 | -0.35 |
|  | Female | 175 | 15.12 | 3.04 |  |  |
| Procedural Knowledge | Male | 175 | 15.89 | 2.88 | 348 | -0.32 |
|  | Female | 175 | 16.02 | 2.85 |  |  |
| Strategic Competence | Male | 175 | 13.40 | 3.12 | 348 | 0.81 |
|  | Female | 175 | 13.15 | 3.05 |  |  |
| Adaptive Reasoning | Male | 175 | 6.78 | 2.55 | 348 | -0.25 |
|  | Female | 175 | 6.82 | 2.62 |  |  |
| Productive Disposition | Male | 175 | 72.35 | 7.50 | 348 | -0.39 |
|  | Female | 175 | 72.52 | 7.44 |  |  |
| Biology Proficiency | Male | 175 | 77.92 | 10.24 | 348 | -0.52 |
|  | Female | 175 | 78.56 | 10.12 |  |  |
| Biology Performance | Male | 175 | 68.41 | 15.32 | 348 | 2.15 |
|  | Female | 175 | 64.29 | 14.97 |  |  |

### Research Question Two

What factors contribute to higher or lower performance in biology among these students?

Table 2 shows that there is a significant positive correlation between student performance in biology and several contributing factors. Specifically, conceptual understanding (Pearson r = .25, p < .01), procedural knowledge (Pearson r = .20, p < .02), and productive disposition (Pearson r = .30, p < .01) each show strong associations with biology performance, indicating that students who score higher in these areas are more likely to perform well in biology. Strategic competence (Pearson r = .15, p < .05) also has a positive, though marginal, relationship with performance, while adaptive reasoning (Pearson r = .10, p = .09) shows a non-significant correlation, suggesting it may play a lesser role in this context. The distinct correlations among the different skill areas suggest that each factor represents a unique aspect of biology proficiency, reinforcing the multifaceted nature of biology understanding and its implications for performance. The findings underscore the importance of conceptual understanding and productive disposition as primary contributors to academic success in biology for technical college students.

**Table 2.** indicates that factors such as conceptual understanding, procedural knowledge, and productive disposition have significant positive correlations with biology performance.

| Factor | Mean<br>(M) | Standard<br>Deviation<br>(SD) | Pearson<br>r | p-value | Interpretation |
| --- | --- | --- | --- | --- | --- |
| Conceptual Understanding | 14.75 | 3.20 | 0.25 | 0.01 | Significant positive relationship with biology performance |
| Procedural Knowledge | 15.90 | 3.15 | 0.20 | 0.02 | Significant positive relationship with biology performance |
| Strategic Competence | 13.25 | 3.10 | 0.15 | 0.05 | Marginal positive relationship with biology performance |
| Adaptive Reasoning | 6.80 | 2.60 | 0.10 | 0.09 | Non-significant positive relationship |
| **Productive Disposition** | 73.00 | 7.35 | 0.30 | 0.01 | Significant positive relationship with biology performance |

### Research Question Three

How do technical college students’ biology proficiency levels compare across regions or schools in Nigeria?

The results in Table 3 below suggest a statistically significant difference in biology proficiency scores across the five regions or schools, with an ANOVA F-value of 4.56 (p = 0.003). This indicates that at least one region or school has significantly different biology proficiency levels compared to the others. Specifically, students from Region 5 (College E) exhibit the highest mean proficiency score (M = 137.90, SD = 9.90), followed by Region 1 (College A) with a mean of 135.40. Regions 2 and 4 show comparatively lower mean scores, suggesting possible variations in instructional quality, resource availability, or other school-specific factors that may influence biology proficiency. These findings highlight the need for targeted interventions in regions with lower proficiency scores to address potential disparities and promote equitable biology education outcomes across schools.

**Table 3:** biology proficiency scores across the five regions or schools.

| School/Region | N | Mean Biology Proficiency Score (M) | Standard Deviation (SD) | ANOVA F-Value | p-value |
| --- | --- | --- | --- | --- | --- |
| Region 1 (College A) | 70 | 135.40 | 10.25 |  |  |
| Region 2 (College B) | 70 | 128.75 | 9.80 |  |  |
| Region 3 (College C) | 70 | 132.30 | 11.10 |  |  |
| Region 4 (College D) | 70 | 130.15 | 10.05 | 4.56 | 0.003 |
| Region 5 (College E) | 70 | 137.90 | 9.90 |  |  |

The results of this study on the biology proficiency of Nigerian technical college students underscore three main findings: first, that gender differences in biology proficiency are minimal; second, that certain cognitive and dispositional factors contribute significantly to higher performance in biology; and third, that notable differences in proficiency exist across various regions or schools. These findings not only reflect the current state of biology education among technical college students but also align with or diverge from previous research in meaningful ways.

The analysis of gender differences in biology proficiency revealed that, although there were slight variations in mean scores, these differences were not statistically significant. Female students achieved slightly higher mean scores than male students, yet this disparity was not large enough to suggest a meaningful gender effect on biology proficiency. This finding is consistent with a growing body of research indicating that gender differences in science proficiency, particularly in biology, are becoming less pronounced as educational opportunities expand for both genders. For example, a study by Oloyede and Afolabi (2020) on secondary school students in Nigeria also found minimal gender differences in biology performance, attributing this parity to increased gender equity in classroom instruction and access to resources. Similarly, global studies, such as those by Hyde and Linn (2006), have suggested that gender gaps in science and mathematics have diminished considerably in recent decades. However, these findings contrast with earlier studies, like that of Nsofor and Bello (2015), which noted significant gender disparities in biology proficiency in certain regions of Nigeria. The discrepancy between older and recent findings may reflect shifts in cultural attitudes and education policies that now encourage female participation in STEM fields.

The second major finding centers on the cognitive and dispositional factors that positively correlate with biology performance. Conceptual understanding, procedural knowledge, and productive disposition each demonstrated a significant positive relationship with biology proficiency. Students with a stronger grasp of biology concepts, an ability to apply learned procedures, and a positive attitude toward the subject achieved higher proficiency scores. This aligns with the findings of Kola and Sunday (2021), who reported that Nigerian students’ success in biology was strongly linked to their conceptual understanding and interest in the subject. Similar results were observed in studies outside Nigeria, such as a 2019 study by Osborne et al., which found that students’ attitudes and dispositions toward science subjects significantly impacted their performance. Conversely, strategic competence and adaptive reasoning, while positively correlated with performance, showed weaker associations in this study. This differs slightly from studies by Adeyemi et al. (2018), which suggested that problem-solving skills (a component of strategic competence) were essential for academic success in biology. The lower correlations observed for these competencies may reflect specific gaps in technical college students’ exposure to biology application and problem-solving exercises, pointing to potential areas for instructional improvement.

Finally, the study revealed significant differences in biology proficiency across various technical colleges in different regions, with students from certain schools displaying higher mean proficiency scores than others. This variation suggests disparities in access to resources, quality of instruction, and possibly even curriculum implementation across regions. Notably, students in Region 5 outperformed their peers, which may indicate that this region has more effective biology teaching methods or better educational resources. This result is consistent with findings from Adebola and Bassey (2022), who noted that urban schools with better resources and qualified teachers often yield students with higher proficiency in biology. The study’s findings on regional disparities align with broader research on educational inequities in Nigeria, such as findings from Ajayi and Ojo (2019), which highlighted how resource allocation often favours certain regions, contributing to academic performance gaps. However, the present study’s findings contradict earlier work by Obe and Martins (2015), which suggested that regional differences in biology performance were insignificant, attributing homogeneity to standardized educational policies. The contrasting findings suggest that while policy may aim for uniform standards, practical differences in regional educational resources continue to influence student outcomes.

These findings offer important insights for policy and practice. The minimal gender differences observed imply that both male and female students in Nigerian technical colleges are benefiting from equitable access to biology education, but continued attention to gender inclusion is necessary to ensure that this parity is maintained. The strong influence of conceptual understanding and productive disposition on biology performance suggests that educators should emphasize these areas, possibly through more engaging, concept-oriented pedagogy that fosters positive student attitudes. Moreover, the observed regional disparities in biology proficiency highlight the need for targeted interventions to address educational inequalities. Improving access to resources and ensuring quality instruction across all technical colleges, particularly in lower-performing regions, could help bridge the gap in biology proficiency.

In conclusion, the study’s findings contribute to a nuanced understanding of factors influencing biology proficiency in Nigerian technical colleges, showing that while gender disparities are minimal, cognitive and dispositional factors and regional educational conditions significantly impact student outcomes. Future research could explore interventions that enhance conceptual understanding and productive disposition while addressing regional resource disparities to promote equitable academic success across Nigeria.

## CONCLUSION

The findings from this study underscore several key insights into factors influencing biology proficiency among technical college students in Nigeria. First, the analysis revealed notable gender differences in biology performance, suggesting that gender-based strategies may be required to address any disparities and enhance overall student achievement in biology. Second, proficiency levels varied significantly across different schools and regions, indicating that resource availability, school policies, and instructional quality likely play substantial roles in shaping students’ biology outcomes. This points to the necessity of policy adjustments that ensure more equitable distribution of educational resources, particularly in under-resourced areas.

Finally, certain factors such as conceptual understanding, strategic competence, and procedural fluency were shown to significantly impact students’ biology proficiency, aligning with previous research on the importance of cognitive skills in science education. However, while regional and school-based differences were observed, these did not uniformly correlate with biology performance, suggesting that institutional quality and support may vary widely within regions. Future studies could benefit from further examining specific institutional practices that enhance biology learning, as well as exploring additional cognitive and socio-environmental variables that may contribute to student success in biology. These findings provide valuable insights for educators, policymakers, and researchers working toward improved science education outcomes in technical colleges across Nigeria.

## RECOMMENDATIONS

The findings of this study have significant implications for both students and biology educators in technical colleges across Nigeria. It is recommended that educators focus on enhancing biology proficiency by integrating diverse instructional strategies that cater to various learning styles and gender differences. Specifically, training teachers to foster a supportive learning environment that encourages collaboration, critical thinking, and exploration of biological concepts can significantly impact student engagement and achievement. Additionally, students should be encouraged to participate in hands-on laboratory activities and collaborative projects that reinforce their conceptual understanding of biology. This approach can help create a community of practice where students can share ideas, challenge each other’s thinking, and develop a deeper understanding of biological principles. Moreover, future research should explore biology proficiency levels in non-elitist technical colleges, which are prevalent throughout Nigeria. Understanding the challenges faced by these institutions can lead to more targeted interventions and policies aimed at improving biology education across diverse contexts. Ultimately, the present study serves as a valuable baseline for understanding biology proficiency among technical college students in Nigeria, and its findings could inform the development of effective educational practices and policies to enhance science education nationwide.

## ACKNOWLEDGEMENTS

The author thanks the Principals, Biology teachers and students of the colleges that participated in the study.

